# Joint Modeling of Reaction Times and Choice Improves Parameter Identifiability in Reinforcement Learning Models

**DOI:** 10.1101/306720

**Authors:** Ian C. Ballard, Samuel M. McClure

**Author notes:** **Corresponding Author**: Ian Ballard.

## Abstract

**Background:** Reinforcement learning models provide excellent descriptions of learning in multiple species across a variety of tasks. Many researchers are interested in relating parameters of reinforcement learning models to neural measures, psychological variables or experimental manipulations. We demonstrate that parameter identification is difficult because a range of parameter values provide approximately equal quality fits to data. This identification problem has a large impact on power: we show that a researcher who wants to detect a medium sized correlation (*r* = .3) with 80% power between a variable and learning rate must collect 60% more subjects than specified by a typical power analysis in order to account for the noise introduced by model fitting.

**New Method:** We derive a Bayesian optimal model fitting technique that takes advantage of information contained in choices and reaction times to constrain parameter estimates.

**Results:** We show using simulation and empirical data that this method substantially improves the ability to recover learning rates.

**Comparison with Existing Methods:** We compare this method against the use of Bayesian priors. We show in simulations that the combined use of Bayesian priors and reaction times confers the highest parameter identifiability. However, in real data where the priors may have been misspecified, the use of Bayesian priors interferes with the ability of reaction time data to improve parameter identifiability.

**Conclusions:** We present a simple technique that takes advantage of readily available data to substantially improve the quality of inferences that can be drawn from parameters of reinforcement learning models.

**Highlights:**

– Parameters of reinforcement learning models are particularly difficult to estimate
– Incorporating reaction times into model fitting improves parameter identifiability
– Bayesian weighting of choice and reaction times improves the power of analyses assessing learning rate

## 1 Introduction

A fundamental question in neuroscience is how animals and humans make decisions for reward or to avoid punishment. In a large class of such decision making problems, animals must use their past experience to infer which action or stimulus leads to the most desirable outcome. This process is thought to occur via processes, collectively known as *reinforcement learning* (RL), in which learning of stimulus or action values is driven by the difference between expectations and outcomes (Rescorla and Wagner, 1972). This difference is known as the *reward prediction error*. Dopamine neurons have been shown to broadcast reward prediction errors in their ongoing firing rate, shaping the value-coding properties of striatal neurons (Lau and Glimcher, 2008; Schultz, 1997). In addition, the reward prediction error has been extensively applied to the interpretation of striatal and midbrain BOLD responses (D’Ardenne et al., 2008; Garrison et al., 2014; McClure et al., 2003), and value estimates from RL models have been used to characterize ventral medial prefrontal cortex (Bartra et al., 2013). To continue making progress in understanding the neural implementation of reward learning, it is necessary to characterize both intra- and inter-individual variability in reinforcement learning due to experimental or trait differences. The most common way to characterize variability is by reference to the learning rate parameter, which controls the effect of prediction errors on subsequent reward predictions. In this paper, we explain why estimation of learning rate is noisy, illustrate the consequences of this noise on power, and propose a method for using reaction time data to mitigate the problem.

Reinforcement learning models have been important in neuroscience in part because they can be applied to any task in which a subject makes repeated decisions. Tasks from human neuropsychology batteries, such as Wisconsin Card Sorting and Weather Prediction, can be understood in the same framework as behavioral neuroscience tasks such as Pavlovian conditioning. We focus our analysis on a task that captures the key features of all reward learning paradigms: the multi-arm bandit. In a bandit task, different options (e.g., levers, slot machines) are associated with a different, and often varying, probability of reward. Animals must decide which option to select based on their experience. This task can be used in rodents, nonhuman primates and humans to understand how animals arbitrate between *exploitation* of an option with a known reward output and *exploration* of alternative options about which the animal knows less (Cohen et al., 2007). Reinforcement learning models are a useful way to characterize behavior and neural systems involved in this task because they formally separate the contribution of learning (i.e., how subjects estimate average reward of each option) from that of exploration/exploitation (Constantino and Daw, 2015; Daw et al., 2006; Kolling et al., 2012; Shenhav et al., 2013; Walton et al., 2010). In contrast, a model-free analysis of choices cannot easily distinguish these two processes.

The learning rate is a natural way to characterize how learning on the bandit task is influenced by factors such as the experimental manipulations, familiarity of the environment, the speed at which the environment changes or the state of the animal. For instance,Behrens et al showed that people learn more quickly (i.e, higher learning rate) when the environment is volatile, and suggested that the anterior cingulate cortex supports this process (Behrens et al., 2007). Although we focus our analysis on the bandit task, we emphasize that learning rate is a useful way to characterize interindividual variability (Frank et al., 2007; Kaiser et al., 2018) or neural function (Costa et al., 2016; Gläscher and O’Doherty, 2010; Schönberg et al., 2007) on any task that can be modeled with reinforcement learning. Therefore, methods that improve the reliability of parameter estimates can benefit a breadth of neuroscience questions that touch upon how animals change their decisions in response to feedback.

We show that learning rate is difficult to estimate because there is a tradeoff between learning rate and a second parameter, decision noise, that results in a range of parameter settings that give near-identical fits to the data. This problem can be intuited through an example. Imagine observing that a subject chose the left lever, received a reward, and subsequently chose the right lever. With these data, it is impossible to know whether this subject did not learn, or learned but responded noisily. Indeed, parameter identifiability is a general problem for models that describe both a hidden psychological process (e.g., learning) and a noisy decision making process. The extent to which this parameter correlation impacts the reliability of learning rate estimates is unknown. We show that unique aspects of reinforcement learning models make them particularly susceptible to parameter correlation.

We propose a method for ameliorating the parameter identifiability problem by harnessing reaction times to constrain estimates of learning rate. Because reaction times are longer for more difficult decisions, they contain rich information about how subjects encode decision variables. Reaction times have been instrumental for understanding how animals navigate a speed/accuracy tradeoff (Ratcliff and McKoon, 2008; Stone, 1960), as well as how neural circuits make decisions about perceptual information (Brody and Hanks, 2016; Shadlen and Newsome, 1998) and reward (Krajbich and Rangel, 2011). In the domain of reinforcement learning, reaction times have been used to a infer learning rate in a task without value-based choice (Bornstein and Daw, 2012). We derive a method for optimally weighting information from choice and reaction times in order to fit reinforcement learning models. Our method for improving the reliability of learning rate estimates could increase the utility of applying reinforcement learning models to the study of behavior, neural systems, and disease (Maia and Frank, 2011).

## 2 Materials and Methods

We contrast parameter estimation in two tasks: a bandit task and a delay discounting task. Code for all simulations and analyses can be found at https://github.com/iancballard/RL-Tutorials.

### 2.1 Task specification: reinforcement learning

We consider a 2-armed bandit task in which the agent decides between two options that independently vary in their probability of reward. Bandits were initialized with a bad arm (35% chance of reward) and a good arm (65% chance of reward). On each trial, the probability of reward for each arm was updated as independent random walks. Thus, the probability of reward on each arm was changed by a random amount taken from Gaussian distribution, *N*(0,.025), with reflecting upper and lower boundaries at 75% and 25% reward, respectively. This design is used to encourage learning over the course of many trials (Daw et al., 2011).

We simulated trajectories of a reinforcement learning agent through the task. The agent tracked values *V* for each of the bandits *s_i_*. On each trial, the agent updates the value of the chosen bandit:

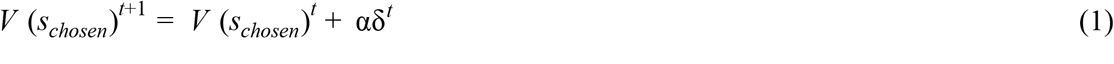

where α is the learning rate and δ^*t*^ is the prediction error from trial *t*:

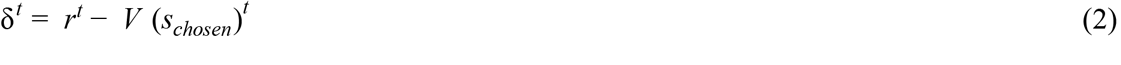

and *r^t^* is the reward received on trial *t*. Values (*V*) were initialized at .5, as there are no negative rewards in this task and this allows for symmetric learning about reward and no-reward outcomes early in the task. Values were transformed into choice probabilities according to a softmax decision rule. If *c^t^* is the choice on trial *t*,

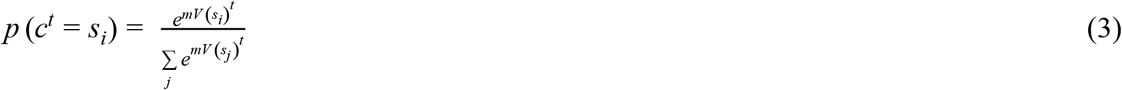

where *m* is the inverse temperature parameter controlling choice variability.

### 2.2 Task specification: delay discounting

Delay discounting requires people to choose between rewards of different amounts (*a*_1_, *a*_2_) available after different delays (*d*_1_, *d*_2_) from the time of the experiment. Hyperbolic discounting describes the subjective value of rewards based on amount and delay:

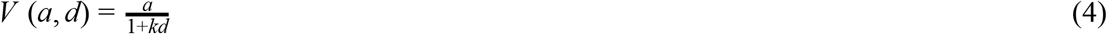

where *k* is the individually determined discount rate that increases with preference for near-term rewards. To create a choice set, we sampled trial-wise *k*s from the same distribution used to generate subject *k*s (see 2.2) and, for each *k*, we set (*a*_1_, *a*_2_, *d*_1_, *d*_2_) to values such that an imaginary subject with that *k* would be indifferent between the two options. This procedure ensures that the choice-set spans the range of indifference points present in the simulated subject population.

Choices in delay discounting are assumed to be described by the softmax function specified in Eqn. 3 except that *V*(*a*_1_,*d*_1_) and *V*(*a*_2_, *d*_2_) replace *V*(*s_j_*). Two parameters therefore determine individual choice behavior in delay discounting: the discount rate, *k*, and the inverse temperature, *m*.

### 2.2 Simulation procedures

Because researchers are often interested in relating learning rates and discount rates to psychological or neural variables of interest, we assessed the extent to which it was possible to recover known parameters from a population of simulated subjects. For each simulation, we drew parameter settings for each subject and generated behavior using the reinforcement learning and delay discounting agents described above. We then fit the parameters of agents to the synthetic data using the Scipy minimize function. We fit the data using four different approaches:

1. Maximum likelihood (ML). This standard technique maximizes the likelihood of the synthetic choice data.
2. ML with reaction times. This technique jointly maximizes the likelihood of the choice and reaction time data.
3. Maximum a posteriori (MAP). This technique uses Bayesian priors to maximize the posterior probability of the choice data.
4. MAP with reaction times. This technique maximizes the posterior probability of the choice and reaction time data.

Finally, we assessed the fitted parameter estimates against the ground truth parameters using Pearson correlation.

We assessed correlations between actual and fitted parameters for each modeling technique and for different numbers of subjects and bandit trials. For set of parameter values, we ran 1,000 simulations. We drew parameters from distributions that matched the expected distribution expected in the population, based on previous literature (Figure S1a, (Daw et al., 2011). A learning rate equal to 0 indicates no learning, whereas a learning rate of 1 indicates a win-stay loose shift strategy. However, low *α* does not necessarily indicate poor performance; rather, it indicates a smaller recency bias in weighting information over past trials (Bayer and Glimcher, 2005). We therefore allowed *α* to span its entire range of values. For delay discounting, we sampled *log(k)* from Ν(− 4.5, .9), Figure S1b (Ballard et al., 2017). For both models, we allowed the inverse temperature to vary from variable (*m* = .5) to nearly deterministic (*m* = 10), Figure S1c. The lower bound on *m* excludes completely noisy subjects; purely random choices are modeled instead as subjects with no learning. Predictions of behavior with α = 0, m > 0 are equivalent to α > 0, m = 0, so we ignore the m = 0 condition without loss of generality. Parameter priors were used to generate simulated subjects and were also used as the Bayesian priors in MAP fitting.

In order to estimate parameter correlations across a range of parameter settings to create Figure 1c,d, we generated simulated data for 1,000 subjects on 200 trials for each cell within a grid of parameter settings. For each simulation run, we numerically computed the Hessian matrix at the maximum likelihood estimate. This matrix describes the convexity of the likelihood function at its peak. We then inverted this matrix to obtain an asymptotic estimate of the variance-covariance matrix. Finally, we converted this matrix to a correlation matrix in order to obtain estimates of the correlation between the parameters at the peak of the likelihood function. Figure 1c,d show the means of these correlations across simulation runs for each parameter setting.

**Figure 1.**
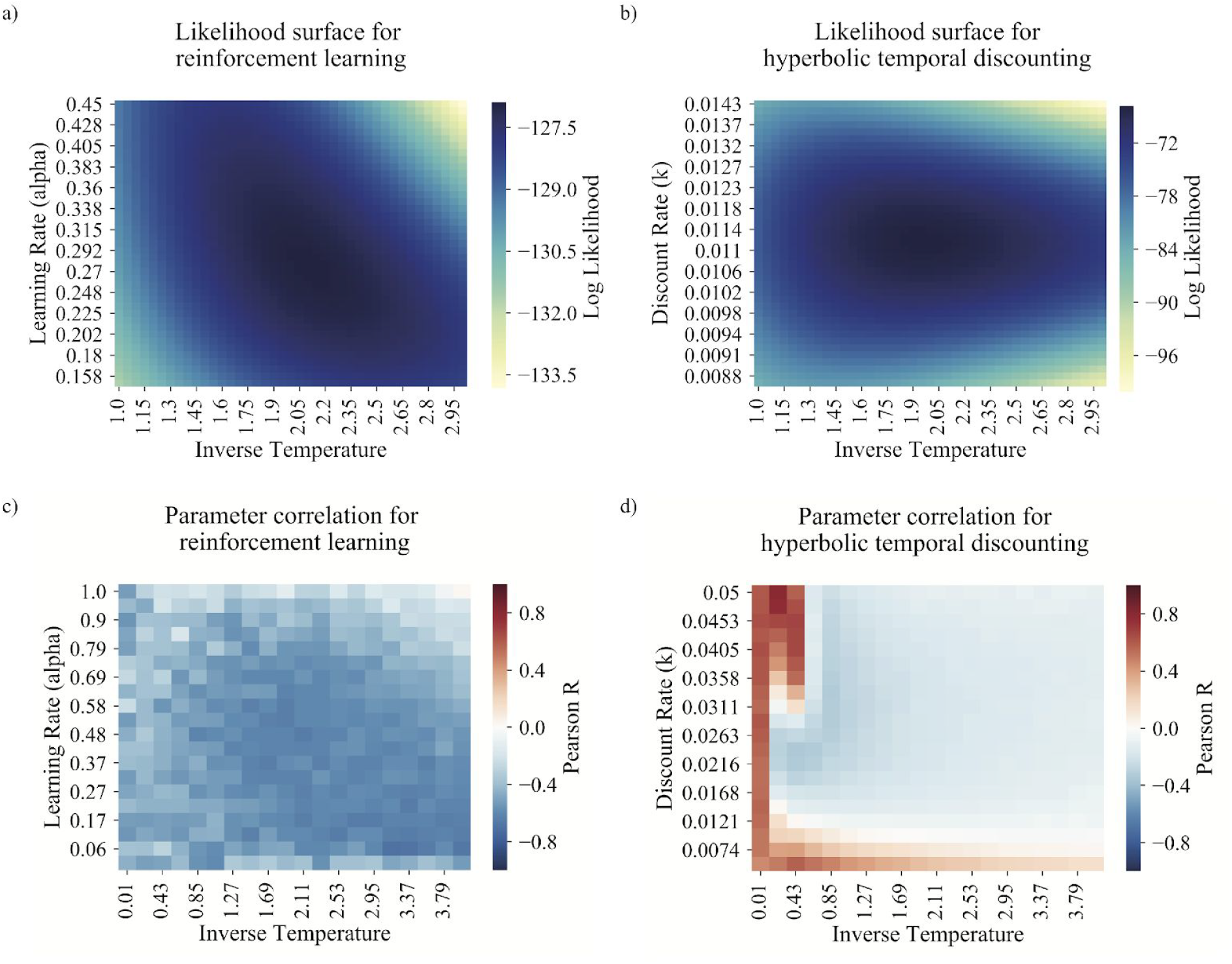
Likelihood surface of reinforcement and delay discounting models. A) Likelihood surface for a reinforcement learning model of a simulated subject on a 2-arm bandit task (*α* = .3, *m* = 2). There is a tradeoff between learning rate and inverse temperature, such that a lower learning rate and more reliable responding provides a similar fit as a higher learning rate and more random responding. B) Likelihood surface for a hyperbolic model of a simulated subject in an delay discounting task (*k* = .011, *m* = 2). Most of the uncertainty comes from the inverse temperature parameter. Compared to A, there is only a modest tradeoff between discount rate and choice noise. C) Estimates of the parameter correlation between *α* and *m* at the maximum likelihood estimate for data simulated from a range of parameter settings. Parameter anticorrelation is high for nearly the full range of parameter settings. D) Estimates of the parameter correlation between *k* and *m* at the maximum likelihood estimate for data simulated from a range of parameter settings. Parameter correlation is high for very noisy subjects and values of *k* near the edges of the choice set, but is generally low for a wide range of typical parameter settings.

### 2.2 Reaction time modeling

We sought to jointly model the probability of the reaction time (*rt*) and choices (*c*). For each trial *t*:

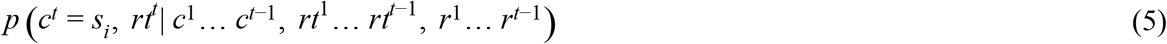

where *r^t^* is the reward earned on trial *t*. Using the chain rule for probabilities, we can rewrite this equation:

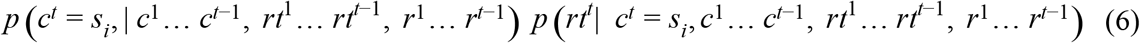

We now make the simplifying assumption that choice on trial *t* is conditionally independent of reaction times on *previous* trials:

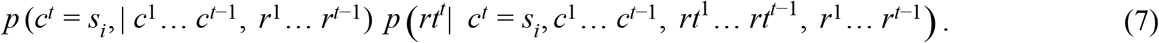

This assumption is sure to be partially invalid because previous trial reaction times are related to previous trial values, which in turn are indirectly related to current trial choice. The complexity of modeling this relationship simultaneously with reinforcement learning is beyond the scope of the current work.

In order to specify a conditional probability distribution on reaction times, we specify a linear regression model relating values of the *p* options as well as the choice to reaction times. We make use of the fact that linear regression can be equivalently specified as a maximum likelihood solution to a linear probabilistic generative model:

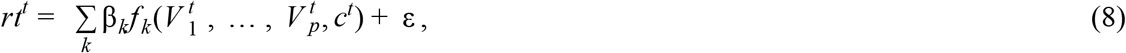

where ε∽ Ν(0, σ^2^). For *n* trials, one can show that the joint log-likelihood of both the observed choices and reaction times (equation 7) is equal to:

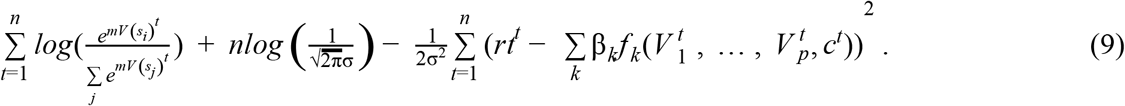

The first term is the softmax log likelihood of choices and the second term is the likelihood of a linear regression model of reaction times, which is included in the output of most linear regression software packages (e.g., statsmodels in Python or logLik in R).

For our simulation data, we created reaction times that were a function of the bandit values. Given the pervasive finding that reaction times are slower for more difficult decisions (Ratcliff and McKoon, 2008), we defined reaction times to be a function of the absolute value of the difference in values between the bandits:

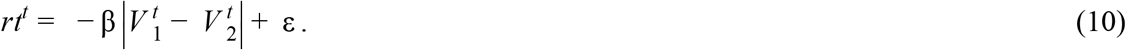

We set β = 1 and added Gaussian random noise to the simulated reaction times with mean of 0 and standard deviation equal to 5 times the standard deviation of 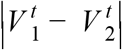 across trials. This procedure resulted in noisy reaction times that were correlated with the absolute value in the difference in choice options with an average *R^2^* = .037. We used a simple model of RTs for the sake of simplicity in simulations. However, one can define a more realistic model and we do so in our analysis of real bandit data.

## 4 Results

We aim to identify and address a problem of identifying the learning rate in RL models. We present a mathematical analysis showing that learning rate is weakly constrained and correlated with decision noise. The specific problem that we highlight is particular to the reinforcement learning algorithm, but parameter identification is an issue for nearly all models. We perform parallel analyses with delay discounting to illustrate the degree to which parameter confusability can differ between different types of models and to show that parameters of reinforcement learning are especially difficult to fit.

### 3.1 Mathematical analysis of parameter identification in reinforcement learning

Parameter identification in reinforcement learning models is encumbered by two sources of ambiguity. First, there is an inherent tradeoff between learning rate and decision noise. Second, the predictions of the model change slowly with changes in learning rate, which exacerbates the trade-off with decision noise. In this section, we present a mathematical analysis illustrating how these two sources of ambiguity arise from the RL algorithm.

We first identify the tradeoff between decision noise and learning rate. Imagine a bandit task to two options: left (s_L_) and right (s_R_). The probability of leftward choice on trial *t* is

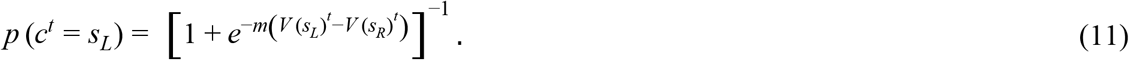

We next examine how this probability changes from trial-to-trial. Imagine that you observe a subject make a leftward choice and receive an outcome. The value of the leftward choice changes according to

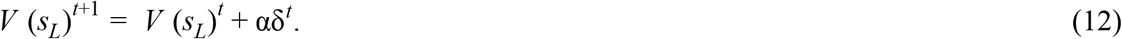

We can insert this update into equation 11 to determine the probability of a leftward choice on the next trial,

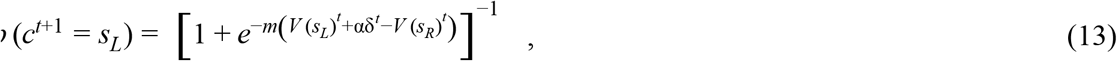

which can be rewritten as

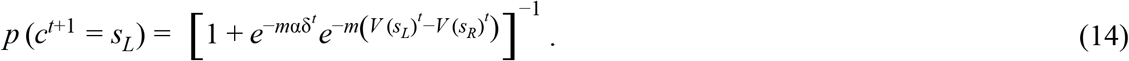

The critical thing to notice is that trial-to-trial changes in behavior (equation 11 compared with equation 14) depends on the product of *m*×*α*. This product indicates an exchangeability between parameters: A high precision (large *m*), slow learning (small *α*) subject would therefore make a similar proportional change in behavior on the subsequent trial as a low precision (small *m*), fast learning (large *α*) subject.

The parameters are not completely redundant because they are constrained by both the prediction error (*δ^t^*), which is a function of the value of the chosen option, and the difference in values, *V* (*s_L_*)^*t*^ − *V* (*s_R_*)^*t*^. These values are a function of both the learning rate and all past outcomes. In principle, this dependence on past outcomes should provide additional information to resolve the learning rate parameter. However, we show that this constraint on learning rate is weak. To do so, we directly compute the value of an option. If *V*(*s_i_*)^0^ is the initial value for action *s_i_* then

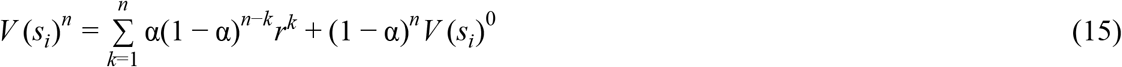

where *n* indexes trials in which *s^i^* is chosen (and hence differs from *t*, used above). The key feature of this equation is that previous trials have an exponentially decreasing influence on value (i.e. (1-*α*)^*n*-*k*^), and the learning rate controls the slope of this exponential decline. Put another way, different learning rates influence value by adjusting the extent to which distant trials influence choice, but these distant trials *always* exert a smaller effect than the most recent trial. As a result, the effect of changing the learning rate on values can be very small. Because values are vital for resolving the ambiguity between learning rate and decision noise, we expect substantial degree of redundancy between these parameters.

### 3.2 Visualizing the scale of parameter redundancy in reinforcement learning

The preceding analysis suggests that the learning rate is poorly constrained, and thus will be correlated with decision noise. However, the magnitude of this correlation is unknown. To estimate the scale of the problem, we plotted the likelihood of the simulated data for all possible parameter settings (i.e., a likelihood surface). Parameter identification involves finding the peak of this likelihood surface. In an ideal setting, the surface would have a single peak with equal curvature in all directions. Parameter correlation manifests as reduced curvature along one direction, indicating that a change of parameters along that direction has a smaller impact on the model fit. Figure 1A shows the likelihood surface for a reinforcement learning model of a simulated subject (*α* = .3, *m* = 2) in a two-arm bandit task. The dark blue area shows a strong tradeoff between parameters; a lower learning rate and reliable responding or higher learning rate and noisier responding provide similar accounts of the data. This flattening of the likelihood surface means than noise in the data can have an outsized effect on the estimates of the peak. If data were only a little different, the peak could move relatively large distances along the flattened direction.

The previous analysis showed parameter correlation for a specific parameter setting. We next assessed whether parameter correlation existed across a range of parameters. We computed the parameter correlation between learning rate and decision noise at the peak of the likelihood surface across a range of parameter settings (See Methods). Figure 1C shows that this tradeoff exists over a wide range of parameters: the average absolute correlation between *α* and *m* is .49. These results suggest that estimates of learning rate are poorly constrained and will be extremely sensitive to noise in the choice data.

### 3.3 Comparison to parameter estimation in delay discounting

Many psychological models separately model a latent psychological process and a noisy decision-making output. An important question is whether all such models suffer from a similar degree of parameter correlation. To begin to address this question, we investigated parameter estimation in delay discounting, a prominent decision making model in neuroscience (McClure et al., 2004). In the prototypical delay discounting task, subjects make a binary choice between a sooner reward, of amount *a_s_* available at delay *d_s_*, and a later reward, of amount *a_ℓ_* and delay *d_ℓ_*. We assumed that people discount delayed rewards according to the hyperbolic discounting model (Eqn. 4) and that choices are made using the softmax decision function (Eqn. 3). The hyperbolic discounting model is parameterized by a a discount rate, *k*, which controls how steeply subjects discount delayed rewards. We first applied a mathematical analysis to assess the degree to which discount rate and decision noise are related. According to the hyperbolic discounting model, the probability of choosing the smaller reward is

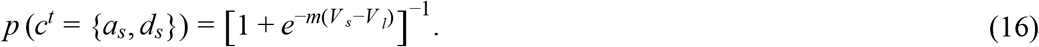

The exponent in this equation is equal to

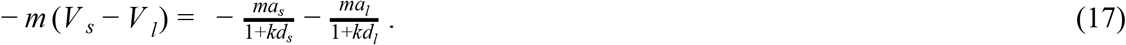

The relationship between the parameters of the model (*k* and *m*) and choice is a function of the amount and delay of the rewards. Because amounts and delays change from trial to trial, the parameters are not generally exchangeable. However, it is possible to construct a scenario in which they are. When *kd_s_*≫1 and *kd_ℓ_*≫1 then (17) is approximated by

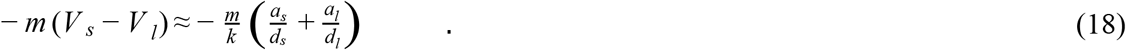

In this regime, *m* and *k* are exchangeable. However, for every delay discounting task that we are aware of, the approximation in Eqn. 18 will not hold for the vast majority of trials. This is because *k* is at most on the order of 10^−2^ (units of day^−1^), intertemporal choice task typically include immediate rewards (*d_s_*= 0), and the longest delays are on the order of one year (10^2^ days). It is therefore generally not the case that *m* and *k* are interchangeable when fitting to delay discounting data.

Based on our mathematical analysis, we predicted that the likelihood surface for hyperbolic discounting would show less parameter correlation than for reinforcement learning. This is evident in Figure 1B. The inverse temperature is not well constrained in our task, but there is relatively little correlation between *m* and *k* in log-likelihood isosurfaces. We quantify this in Figure 1D, which shows that, over a range of parameter settings, the average absolute correlation between *m* and *k* is .19. Notably, this correlation mostly arises at extreme values of the parameters that are unlikely to occur in many subjects. This example illustrates that reinforcement learning is much more susceptible to parameter identifiability problems than delay discounting. More generally, we can expect some degree of parameter tradeoff in many psychological models, but the features of RL models we identify in *3.1* make the parameters of RL models particularly difficult to estimate.

### 3.4 Reaction times and Bayesian priors improve parameter identifiability

The previous sections demonstrate that learning rate is poorly constrained in typical experimental settings. A consequence of this is that its estimation will be sensitive to idiosyncrasies in the choice data. However, the scale of this problem is unknown. To address this question, we asked whether we could recover learning rates from a cohort of simulated subjects. This question is important because many experiments relate learning rate to a neural or behavioral variable of interest (Behrens et al., 2007; Rouhani et al., 2018) and errors in estimates of learning rate reduce the power of these tests. For each subject in a cohort, we simulated choices on a two-arm bandit using the RL model described. We then fit this simulated data with the same RL model and computed the correlation between the ground truth learning rates and the recovered learning rates. Higher correlations indicate a better ability to estimate parameters for a group of subjects. As anticipated by the previous results, the ability to reconstruct discount rates in delay discounting is near ceiling (Figure S2). We therefore focus our analyses on parameter identifiability in RL.

We find that the ability to reconstruct learning rates is modest. The correlation between the model-fit parameters and the ground truth values was, for 100 trials of data, *r* = .66, and for 200 trials, *r* = .79. This number of trials is typical in bandit tasks (Wimmer et al., 2014; Wunderlich et al., 2009). Note that these simulations are likely to overestimate the true ability to reconstruct learning rates because the subjects were simulated and fit with the same learning model. In real data, RL models only approximate all the psychological processes at play. Therefore, these correlation estimates can be viewed as an approximate upper bound on the ability to fit learning rates.

We computed examined parameter identifiability as a function of the number of trials and the number of subjects in a cohort (Figure 2). We found that increasing the number of trials improves the ability to reconstruct learning rates. If there are more trials, learning rate will be constrained by a richer set of choices with different recent reward histories between the two options. We also found that, perhaps surprisingly, increasing the number of subjects does not appreciably improve the ability to reconstruct learning rates. Because the noise in parameter fitting is at the individual subject level, larger sample sizes do not improve parameter identifiability. Larger sample sizes will always increase statistical power for group tests; however, they are not a solution to the parameter identifiability problem in reinforcement learning.

**Figure 2.**
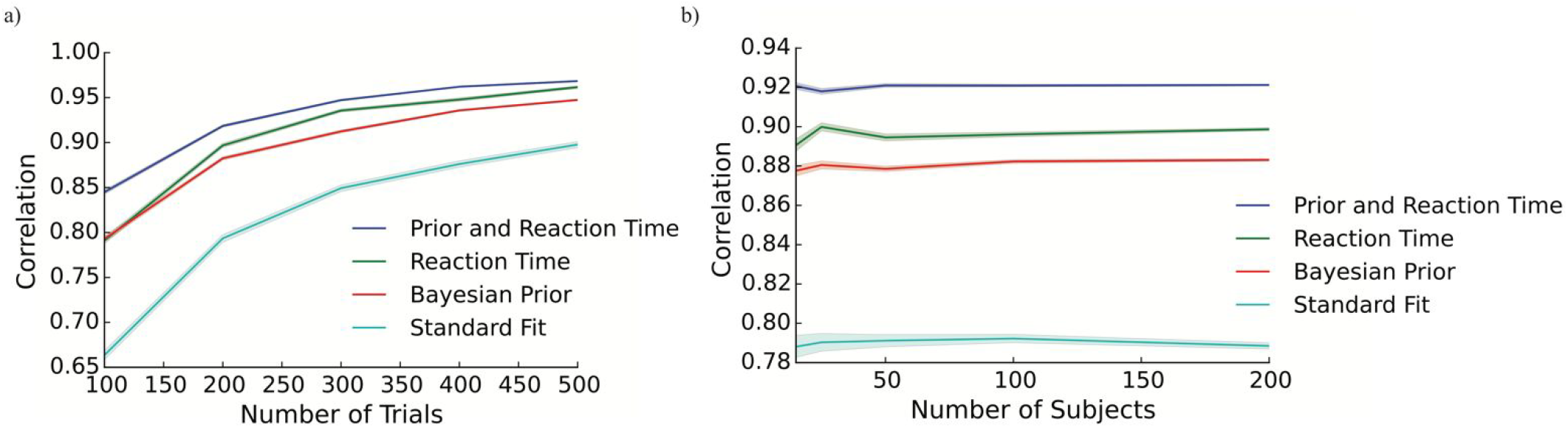
Simulation of parameter identifiability in RL. A) The correlation between ground truth and fitted learning rates as a function of the number of bandit trials (25 subjects). Increasing the number of trials improves parameter identifiability, and the use of reaction times and Bayesian priors substantially improves parameter identifiability regardless of the number of trials. B) The correlation between ground truth and fitted learning rates as a function of the number of subjects (200 trials). Increasing the number of subjects did not improve parameter identifiability. The use of reaction times and Bayesian priors substantially improves parameter identifiability regardless of the number of subjects.

We propose using reaction times as an additional source of information to constrain learning rate and improve parameter identifiability. Our method jointly maximizes the likelihood of both the choice and the reaction time data (See Methods). We assessed the efficacy of this technique against a different approach for improving parameter identifiability: use of Bayesian priors (Gershman, 2016). Bayesian priors can prevent overfitting by rendering certain parameter values unlikely, which reduces the tradeoff between correlated parameters. We also tested whether the use of both reaction times and Bayesian priors could have an additive effect on parameter identifiability (Figure 2B). All tests were Bonferroni corrected for 10 tests across the trial and subject bins.

The use of either reaction times or Bayesian priors improved parameter identifiability for all trial counts and all cohort sizes, all *p* < .001. Reaction times improved parameter identifiability more than Bayesian priors, 15 subjects/100 trials, *p* = .019, all others, *p* < .001, except for bandits with 100 trials, *p* > .2. This finding is remarkable given that reaction times were noisily related to the difference in option values (average *R^2^* = .037), and therefore contained limited information about learning rate. Further, the Bayesian prior approach benefited from the fact that the priors were perfectly accurate: simulated subject parameters were drawn from the same prior distributions used for Bayesian model fitting. Finally, we found that the combined use of Bayesian priors and reaction times provided the best fit for all cohort sizes and numbers of trials, all *p* < .001. This finding confirms that reaction times and Bayesian priors offer partially distinct ways to regularize RL parameter estimates, and their combined use further improves estimates.

### 3.5 Effect of parameter identifiability on experimental power

Our analysis of the ability to recover the learning rates of a subject population are optimistic because we simulated and fit data with the same model. Even so, the simulated correlation between recovered learning rates and ground truth may appear high. However, even moderate decrements in the ability recover learning rates can have a large effect on experimental power (Figure 3). For example, one needs around 85 subjects to detect a correlation of *r* = .3 with 80% power. If experimental constraints set the maximum number of trials to be 200, then our simulations suggest that experimenter would need to collect data from *at least* 137 subjects to detect the same effect with the same power, a 60% increase. According to our simulations, the use of reaction times could increase parameter identifiability sufficiently to require 105 subjects. The use of these model-fitting techniques can appreciably improve power.

**Figure 3.**
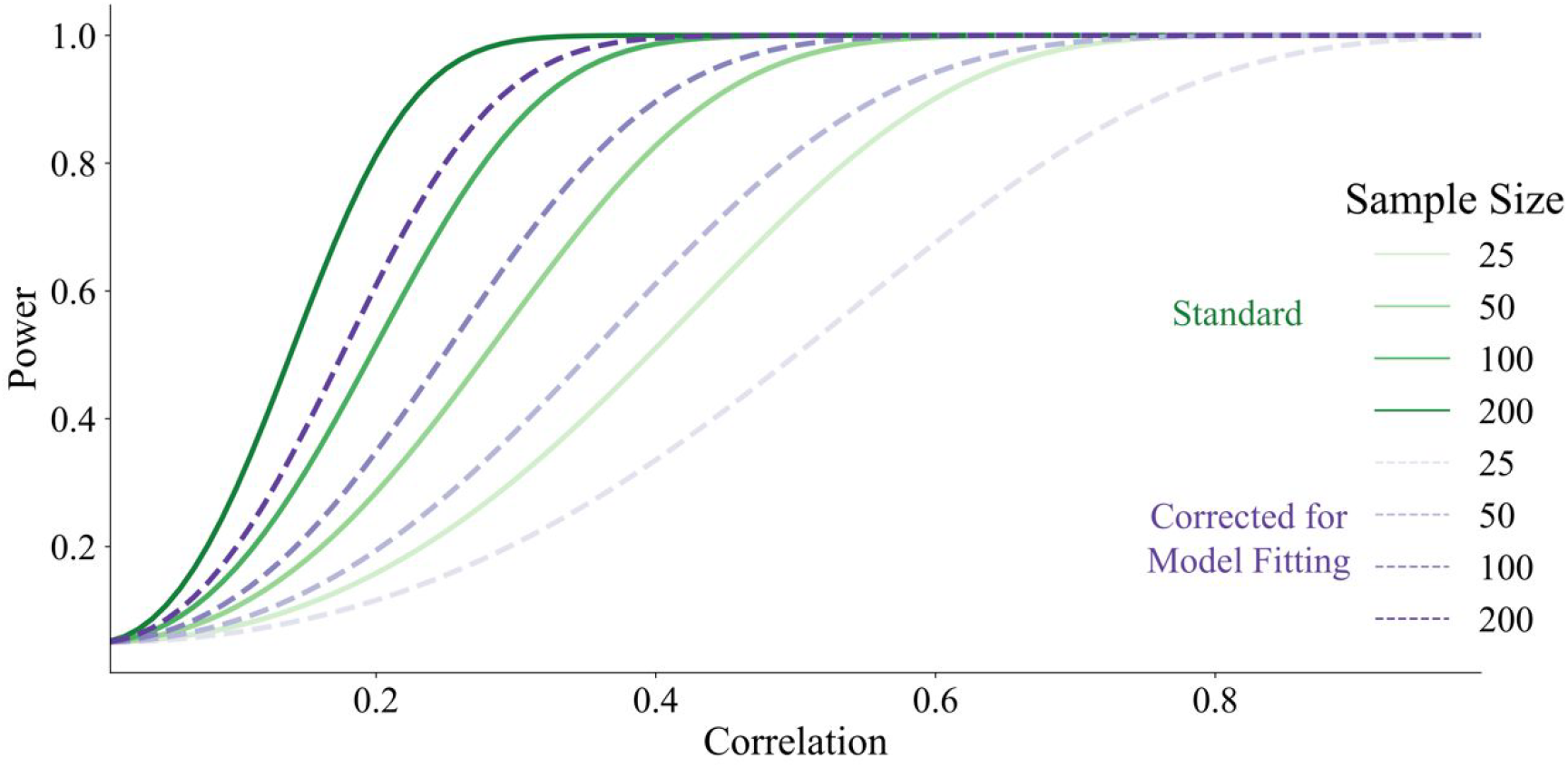
Effect of parameter identifiability on experimental power. Green lines depict the power to detect a correlation between two variables for different sample sizes. Purple lines depict the power after accounting for the noise introduced by model-fitting.

### 3.6 Application to Empirical Data

Our simulations showed that both the use of a Bayesian prior and the use of reaction times in model fitting improved parameter identifiability. Although these simulations were designed to match experimental data as closely as possible, they included two important features that could render their predictions over-optimistic. First, the parameters were generated from the same prior distribution used for fitting with Bayesian priors. Second, the reaction times were generated from the model used to fit reaction times. In order to assess the efficacy of the proposed method in a more realistic setting, we turned to a published dataset (Wimmer et al., 2014). Thirty subjects performed 100 trials of a two-armed bandit task in which each bandit was associated with a different, trial-unique object that was irrelevant to the task. In addition, subjects performed a second run of the task without the objects. Although there were differences between the tasks (e.g., presence of objects), and there could have been state differences within a subject (e.g., fatigue), we reasoned that learning rates assessed from two very similar bandit tasks should be strongly correlated. We modeled log transformed reaction times as a function of 1) the linear and quadratic effects of the absolute value of the difference in values between the bandits 2) the linear effect of trial number, and an 3) indicator function on whether the subject choose the bandit with the maximum value:

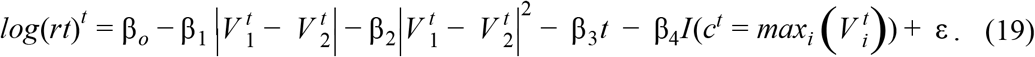

Where (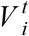) is the index of the maximum value and *I* is an indicator function.

Under standard model fitting with no Bayesian priors and no reaction times, learning rates are correlated, *r*(28) = .47, *p* = .009, Figure 4A. Fitting with Bayesian priors results in a larger correlation, *r*(28) = .59. We used the R package Bayesian First Aid to assess the difference in correlations and found a 74.5% posterior probability that Bayesian priors improved the correlation. Fitting with reaction times results in an even higher correlation, *r*(28) = .75, Figure 4B. The Bayesian posterior probability that reaction times increased the magnitude of the correlation over standard model fitting is 94.5%. In contrast to our simulation results, the combined use of both Bayesian priors and RT results in a similar strength correlation as using only Bayesian priors, *r*(28) = .60, *p* < .001. One possibile account for this finding is that reaction time modeling has no effect over-and-above Bayesian priors. However, the parameters from combined reaction times and Bayesian fitting were only moderately correlated with the parameters from using Bayesian priors, *r*(28) = .62, *p* < .001, or reaction times, *r*(28) = .55, *p* =.002, alone. Therefore, the use of Bayesian priors worsened the fit relative to using reaction times alone. We speculate that this is because our Bayesian priors did not perfectly align with the distribution of parameters in our subjects, and therefore reaction times and Bayesian priors have different effects on parameter estimates. Overall, our results suggests that fitting reaction times improves the ability to estimate learning rates in empirical bandit data.

**Figure 4.**
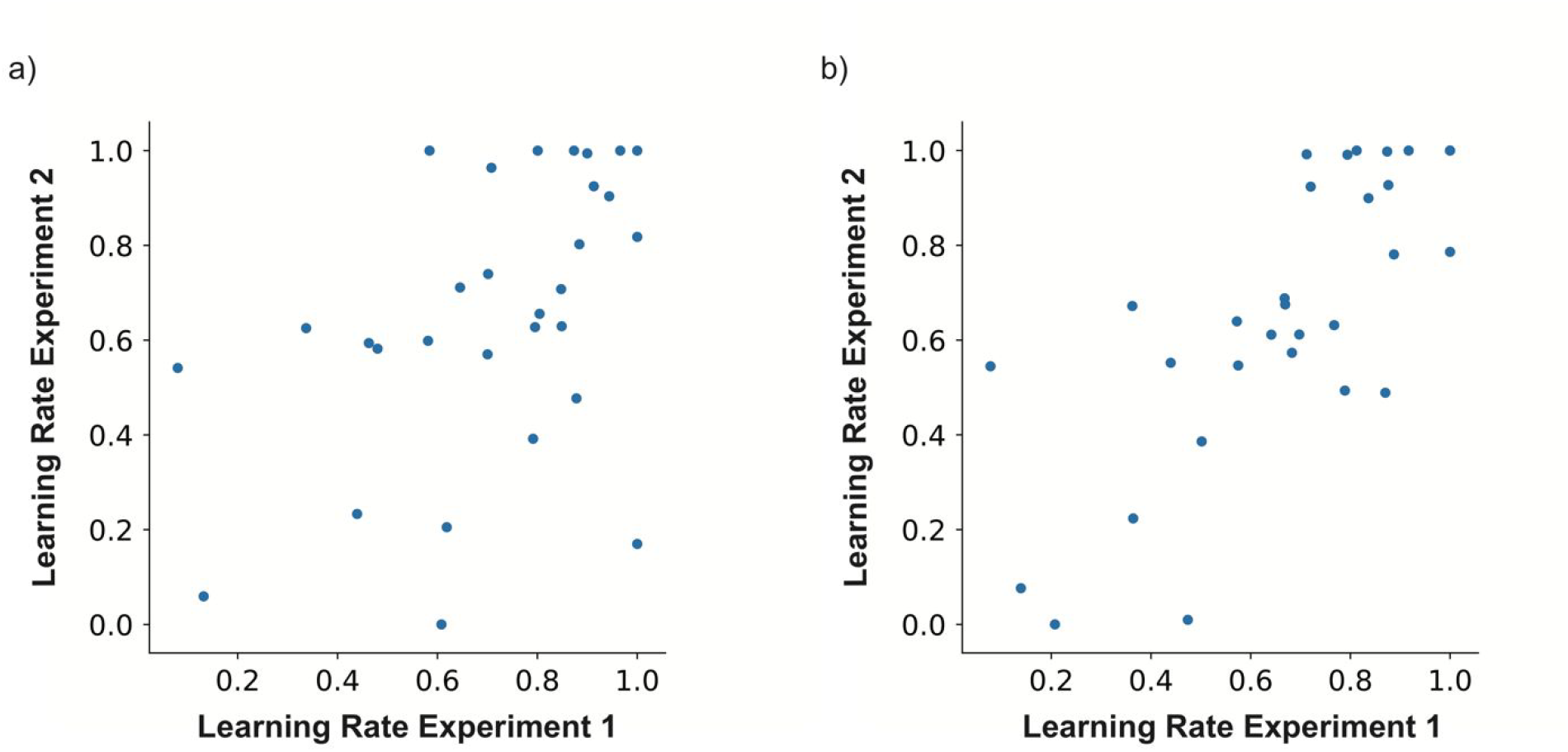
Joint modeling of choice and reaction times improves parameter identifiability in real data. A) The correlation in estimated learning rate between two runs of a bandit task using a standard RL model. B) The correlation in estimated learning rate when estimated using a model of reaction times and choice.

## 4 DISCUSSION

The utility of reinforcement learning models depends on the experimenter’s ability to estimate its parameters. We showed that jointly fitting choices and reactions times improves the reliability of RL parameter estimates. Some researchers have used Bayesian priors in order to improve the quality of reinforcement learning model fits (Daw et al., 2011). We found that Bayesian priors actually interfered with the ability of reaction times to improve parameter identifiability in real data, potentially because the priors were misspecified (section *3.6*). A major advantage of our method is that fitting based on reaction times requires no prior knowledge of the subject population. More sophisticated Bayesian approaches share this advantage. Recent methods derive empirical priors from an independent dataset (Gershman, 2016), or hierarchically fit group-level priors that constrain subject-level parameters (Chávez et al., 2017). However, the efficacy of these approaches is contested (Spektor and Kellen, 2018) and they can yield counterintuitive results. For instance, the empirical prior on learning rate advocated by Gershman is concentrated entirely on 0 and 1. This prior expresses the belief that subjects do not use reinforcement learning: they either don’t learn, or they do use a win-stay/lose-shift strategy. Although this prior is empirically derived, it expresses a belief about learning that is opposed by a wealth of behavioral and neural data. Because our method capitalizes on new information available in reaction times, we argue that it is less susceptible to noise in the choice data than methods that use choice data to specify Bayesian priors. Nonetheless, we did not implement these more sophisticated Bayesian approaches and it is possible that a hybrid approach using both empirical priors and reaction times confers the largest benefit on parameter identifiability.

There is a growing appreciation of the importance of statistical power and the prevalence of underpowered studies in experimental neuroscience. Low power is a twofold blow to experiments: it both reduces the likelihood of detecting an effect and reduces the probability that an effect, once detected, is true (Button et al., 2013). Investigations of across-subject variability in learning rate must take into account that power for across-subject tests is lower than for between-subject tests (Yarkoni, 2009). We have shown that the noise introduced by model-fitting appreciably reduces the power of a between-subjects correlation. Notably, the reduction in power is larger for smaller samples. As a result, already underpowered studies will be especially prone to Type I and Type II errors when analyzing across-subject relationships with learning rate. While our method ameliorates this noise, it does not eliminate it. We recommend that the simulations in Figure 2 inform power analyses for across-subject studies. However, we emphasize that the estimates in Figure 2 are likely over-optimistic for real data. Finally, our results show that that increasing the number of subjects will increase power (Figure 3), but does not reduce the effect of noise on parameter identification (Figure 2). In contrast, increasing the number trials ameliorates the effect of noise and increases power.

Our modeling approach focused on a simple reinforcement learning model applied to a bandit task. There is substantial interest in investigating parameters of more complex RL tasks. For example, in the two-step decision task, subjects make sequential decisions to earn rewards (Daw et al., 2011). This behavior is well-described by a hybrid between a model-free learning agent, similar to the one described here, and a model-based agent that makes decisions based on a model of the sequential structure of the task. Experimenters have estimated the relative weighting between these two systems and have related the weighting parameter to individual differences in neural activation (Daw et al., 2011; Doll et al., 2015), working memory capacity (Otto et al., 2013), habit persistence (Gillan et al., 2015) and compulsive behavior (Gillan et al., 2016). Because hybrid models depend on the same reinforcement learning mechanism described here, it is likely that at least some of the parameters in these models suffer from similar fitting problems. We expect that joint modeling of reaction times could help improve estimates of individual differences in goal-directed behavior.

Formal models are vital for developing a theoretical understanding of brain and behavior (Gläscher and O’Doherty, 2010). However, most models contain hidden complexity that can reduce their applicability. In RL, this complexity is due to the fact that the behavior of the model changes slowly with changes in parameterization (Wilson and Niv, 2015). Faced with this problem, experimenters should make use of all the information available to help constrain estimates of learning. Reaction times are a useful and readily available source of such information. Future work should consider how biometrics such as eye tracking could be used to further constrain model estimates (Leong et al., 2017).

## 5 Acknowledgements

The authors would like to thank Elliott Wimmer for providing data, Yuan Chang Leong for feedback and the NSF GRFP and NSF IGERT grant for providing training support for I.B.

**Author Contributions**: IB conceptualized the study, conducted the analysis, and wrote the manuscript. SM provided supervision and critical revisions.
Abstract

## References

Ballard, I.C., Kim, B., Liatsis, A., Aydogan, G., Cohen, J.D., McClure, S.M., 2017. More Is Meaningful: The Magnitude Effect in Intertemporal Choice Depends on Self-Control. Psychol. Sci. 27, 956797617711455–956797617711454.

Bartra, O., McGuire, J.T., Kable, J.W., 2013. The valuation system: A coordinate-based meta-analysis of BOLD fMRI experiments examining neural correlates of subjective value. Neuroimag. 76, 412–427.

Bayer, H.M., Glimcher, P.W., 2005. Midbrain dopamine neurons encode a quantitative reward prediction error signal. Neuro. 47, 129–141.

Behrens, T.E.J., Woolrich, M.W., Walton, M.E., Rushworth, M.F.S., 2007. Learning the value of information in an uncertain world. Nature Publishing Grou. 10, 1214.

Bornstein, A.M., Daw, N.D., 2012. Dissociating hippocampal and striatal contributions to sequential prediction learning. Eur. J. Neurosci. 35, 1011–1023.

Brody, C.D., Hanks, T.D., 2016. Neural underpinnings of the evidence accumulator. Curr. Opin. Neurobiol. 37, 149–157.

Button, K.S., Ioannidis, J.P.A., Mokrysz, C., 2013. Power failure: why small sample size undermines the reliability of neuroscience. … Reviews Neuroscience.

Chávez, M.E., Villalobos, E., Baroja, J.L., Bouzas, A., 2017. Hierarchical Bayesian modeling of intertemporal choice. Judgement and Decision Makin. 12, 19–28.

Cohen, J.D., McClure, S.M., Yu, A.J., 2007. Should I stay or should I go? How the human brain manages the trade-off between exploitation and exploration. Philos. Trans. R. Soc. Lond. B Biol. Sci. 362, 933–942. https://doi.org/10.1098/rstb.2007.2098

Constantino, S.M., Daw, N.D., 2015. Learning the opportunity cost of time in a patch-foraging task. Cogn. Affect. Behav. Neurosci. 15, 837–853. https://doi.org/10.3758/s13415-015-0350-y

Costa, V.D., Dal Monte, O., Lucas, D.R., Murray, E.A., Averbeck, B.B., 2016. Amygdala and Ventral Striatum Make Distinct Contributions to Reinforcement Learning. Neuro. 92, 505–517.

D’Ardenne, K., McClure, S.M., Nystrom, L.E., Cohen, J.D., 2008. BOLD responses reflecting dopaminergic signals in the human ventral tegmental area. Scienc. 319, 1264–1267.

Daw, N.D., Gershman, S.J., Seymour, B., Dayan, P., Dolan, R.J., 2011. Model-based influences on humans’ choices and striatal prediction errors. Neuro. 69, 1204–1215.

Daw, N.D., O’Doherty, J.P., Dayan, P., Seymour, B., Dolan, R.J., 2006. Cortical substrates for exploratory decisions in humans. Natur. 441, 876–879.

Doll, B.B., Duncan, K.D., Simon, D.A., Shohamy, D., Daw, N.D., 2015. Model-based choices involve prospective neural activity. Nat. Neurosci. 18, 767–772.

Frank, M.J., Moustafa, A.A., Haughey, H.M., Curran, T., Hutchison, K.E., 2007. Genetic triple dissociation reveals multiple roles for dopamine in reinforcement learning. Proceedings of the National Academy of Science. 104, 16311–16316.

Garrison, J., Erdeniz, B., Done, J., 2014. Corrigendum to “Prediction error in reinforcement learning: A meta-analysis of neuroimaging studies” [Neurosci. Biobehav. Rev. 37 (7), (2013) 1297–1310]. Neurosci. Biobehav. Rev. 47, 754. https://doi.org/10.1016/j.neubiorev.2014.10.010

Gershman, S.J., 2016. Empirical priors for reinforcement learning models. J. Math. Psychol. 71, 1–6.

Gillan, C.M., Kosinski, M., Whelan, R., Phelps, E.A., Daw, N.D., 2016. Characterizing a psychiatric symptom dimension related to deficits in goal-directed control. Elif. 5, e94778.

Gillan, C.M., Otto, A.R., Phelps, E.A., Daw, N.D., 2015. Model-based learning protects against forming habits. Cogn. Affect. Behav. Neurosci. 15, 523–536.

Gläscher, J.P., O’Doherty, J.P., 2010. Model-based approaches to neuroimaging: combining reinforcement learning theory with fMRI data. Wiley Interdiscip. Rev. Cogn. Sci. 1, 501–510.

Kaiser, R.H., Treadway, M.T., Wooten, D.W., Kumar, P., Goer, F., Murray, L., Beltzer, M., Pechtel, P., Whitton, A., Cohen, A.L., Alpert, N.M., El Fakhri, G., Normandin, M.D., Pizzagalli, D.A., 2018. Frontostriatal and Dopamine Markers of Individual Differences in Reinforcement Learning: A Multi-modal Investigation. Cereb. Corte. 28, 4281–4290. https://doi.org/10.1093/cercor/bhx281

Kolling, N., Behrens, T.E.J., Mars, R.B., Rushworth, M.F.S., 2012. Neural Mechanisms of Foraging. Scienc. 336, 95–98. https://doi.org/10.1126/science.1216930

Krajbich, I., Rangel, A., 2011. Multialternative drift-diffusion model predicts the relationship between visual fixations and choice in value-based decisions. Proc. Natl. Acad. Sci. U. S. A. 108, 13852–13857. https://doi.org/10.1073/pnas.1101328108

Lau, B., Glimcher, P.W., 2008. Value Representations in the Primate Striatum during Matching Behavior. Neuro. 58, 451–463.

Leong, Y.C., Radulescu, A., Daniel, R., DeWoskin, V., Niv, Y., 2017. Dynamic Interaction between Reinforcement Learning and Attention in Multidimensional Environments. Neuro. 93, 451–463.

Maia, T.V., Frank, M.J., 2011. From reinforcement learning models to psychiatric and neurological disorders. Nature Publishing Grou. 14, 154–162.

McClure, S.M., Berns, G.S., Montague, P.R., 2003. Temporal prediction errors in a passive learning task activate human striatum. Neuro. 38, 339–346.

McClure, S.M., Laibson, D.I., Loewenstein, G., Cohen, J.D., 2004. Separate neural systems value immediate and delayed monetary rewards. Scienc. 306, 503–507. https://doi.org/10.1126/science.1100907

Otto, A.R., Raio, C.M., Chiang, A., Phelps, E.A., Daw, N.D., 2013. Working-memory capacity protects model-based learning from stress. Proc. Natl. Acad. Sci. U. S. A. 110, 20941–20946.

Ratcliff, R., McKoon, G., 2008. The Diffusion Decision Model: Theory and Data for Two-Choice Decision Tasks. dx.doi.org.stanford.idm.oclc.or. 20, 873–922.

Rescorla, R., Wagner, A., 1972. A Theory of Pavlovian Conditioning: Variations in the Effectiveness of Reinforcement and Nonreinforcement (Appletone-Century-Crofts, New York).

Rouhani, N., Norman, K.A., Niv, Y., 2018. Dissociable effects of surprising rewards on learning and memory. J. Exp. Psychol. Learn. Mem. Cogn.

Schönberg, T., Daw, N.D., Joel, D., O’Doherty, J.P., 2007. Reinforcement learning signals in the human striatum distinguish learners from nonlearners during reward-based decision making. J. Neurosci. 27, 12860–12867.

Schultz, W., 1997. A Neural Substrate of Prediction and Reward. Scienc. 275, 1593–1599.

Shadlen, M.N., Newsome, W.T., 1998. The variable discharge of cortical neurons: implications for connectivity, computation, and information coding. J. Neurosci. 18, 3870–3896.

Shenhav, A., Botvinick, M.M., Cohen, J.D., 2013. The expected value of control: an integrative theory of anterior cingulate cortex function. Neuro. 79, 217–240.

Spektor, M.S., Kellen, D., 2018. The relative merit of empirical priors in non-identifiable and sloppy models: Applications to models of learning and decision-making: Empirical priors. Psychon. Bull. Rev. 1–22.

Stone, M., 1960. Models for choice-reaction time. Psychometrik. 25, 251–260.

Walton, M.E., Behrens, T.E.J., Buckley, M.J., Rudebeck, P.H., Rushworth, M.F.S., 2010. Separable Learning Systems in the Macaque Brain and the Role of Orbitofrontal Cortex in Contingent Learning. Neuro. 65, 927–939. https://doi.org/10.1016/j.neuron.2010.02.027

Wilson, R.C., Niv, Y., 2015. Is Model Fitting Necessary for Model-Based fMRI? PLoS Comput. Biol. 11, e1004237–21.

Wimmer, G.E., Braun, E.K., Daw, N.D., Shohamy, D., 2014. Episodic Memory Encoding Interferes with Reward Learning and Decreases Striatal Prediction Errors. J. Neurosci. 34, 14901–14912.

Wunderlich, K., Rangel, A., O’Doherty, J.P., 2009. Neural computations underlying action-based decision making in the human brain. Proc. Natl. Acad. Sci. U. S. A. 106, 17199–17204.

Yarkoni, T., 2009. Big Correlations in Little Studies: Inflated fMRI Correlations Reflect Low Statistical Power—Commentary on Vul et al. (2009). Perspect. Psychol. Sci. 4, 294–298.

